# Selection of pre-leukemic hematopoietic stem cells driven by distinct extracellular matrix molecules

**DOI:** 10.1101/2025.01.20.633881

**Authors:** Maria Jassinskaja, Daniel Bode, Monika Gonka, Theodoros I Roumeliotis, Alexander J Hogg, Juan A Rubio Lara, Ellie Bennett, Joanna Milek, Bart Theeuwes, M S Vijayabaskar, Lilia Cabrera Cosme, James L C Che, Sandy MacDonald, Sophia Ahmed, Benjamin A Hall, Grace Vasey, Helena Kooi, Miriam Belmonte, Mairi S Shepherd, William J Brackenbury, Iwo Kucinski, Satoshi Yamazaki, Andrew N Holding, Alyssa H Cull, Nicola K Wilson, Berthold Göttgens, Jyoti Choudhary, David G Kent

## Abstract

Despite rapid advances in mapping genetic drivers and gene expression changes in hematopoietic stem cells (HSCs), there is a relative paucity of studies at the protein level. Here, we perform a deep, multi-omic characterization (epigenome, transcriptome and proteome) of HSCs carrying a loss-of-function mutation in *Tet2*, a key driver of increased self-renewal in blood cancers. Using state-of-the-art, multiplexed, low-input mass spectrometry (MS)-based proteomics, we profile wildtype (WT) and TET2-deficient (*Tet2*^-/-^) HSCs and show that the proteome captures previously unrecognized molecular processes which define the pre-leukemic HSC molecular landscape. Specifically, we obtain more accurate stratification of WT and *Tet2*^-/-^ HSCs than transcriptomic approaches and identify extracellular matrix (ECM) molecules as novel points of dysregulation upon TET2 loss. HSC expansion assays using ECM-functionalized hydrogels confirm a selective effect on the expansion of *Tet2*-mutant HSCs. Taken together, our study represents a comprehensive molecular characterization of *Tet2*-mutant HSCs and identifies a previously unanticipated role of ECM molecules in regulating self-renewal of disease-driving HSCs.

## Introduction

Hematological malignancies are commonly initiated by single hematopoietic stem cells (HSCs) that have acquired mutations which confer a clonal advantage relative to non-mutated hematopoietic cells^1^. Loss-of-function (LOF) mutations in the epigenetic regulator TET2 are commonly found in hematological malignancies and evidence points toward its loss driving an increase in HSC self-renewal^1–5^. At the transcript level, mutations in *Tet2* are associated with altered gene expression in mouse HSCs; however, relatively few of these potential partner genes have been implicated in directly driving disease initiation and progression^2–4^, thus highlighting an urgent need to further explore the molecular landscape of mutated HSCs.

Historically, comprehensive profiling of HSCs beyond the transcriptome has been impeded due to their low numbers. To address the prohibitively large amount of material typically required for global proteomic characterization, multiple strategies for facilitating low cell number and single cell mass spectrometry (MS)-based proteomics have begun to emerge^6–12^ . These efforts have revealed a generally poor correlation between proteome and transcriptome in hematopoietic stem and progenitor cells (HSPCs), especially in non-homeostatic contexts such as inflammation and disease^10,11,13,14^, further highlighting the need to develop low cell number approaches to facilitate study of the global proteome in HSCs and access the functional unit of molecular activity.

In this study, we use an MS-based method for global proteomic profiling of low numbers (10,000-20,000) of primary HSPCs as a part of a comprehensive multi-omic profiling of Tet2-deficient (*Tet2*^-/-^) long term (LT)-HSCs and uncover new regulators of *Tet2* mutant HSC biology. Analysis of the proteome exclusively identifies extracellular matrix (ECM) interactions as a point of dysregulation upon loss of TET2 in HSCs, providing the first evidence for ECM molecules promoting the self-renewal of mutant HSCs relative to their non-mutant counterparts, and thus identifies previously unexplored pathways for therapeutic intervention. These data highlight the importance of assessing the proteome of pre-leukemic and leukemic HSCs in order to reveal novel biology that is typically hidden from genomic and transcriptomic studies.

## Results

### Integrative single cell ATAC-seq and RNA-seq analysis of TET2-deficient HSCs

To understand molecular changes in HSCs induced by TET2 LOF, we first assessed chromatin accessibility and gene expression at the single-cell level. Using a genetic knockout mouse model with targeted disruption of the TET2 catalytic domain^5^, we applied single-cell Assay for Transposase-Accessible Chromatin by sequencing (scATAC-seq; Figures 1A-1C) and single-cell RNA sequencing^15^ (scRNA-seq; Figures 1D and 1E) to fluorescence activated cell sorting (FACS)-isolated CD45^+^ CD48^-^ CD150^+^ EPCR^+^ Sca-1^+^ (ESLAM Sca-1^+^) HSCs, a cell population highly enriched (>60%) for HSCs with long-term (LT) serial reconstitution capacity^16^ from *Tet2*^-/-^ and wild-type (WT) mice. scATAC-seq analysis revealed changes in chromatin accessibility induced by TET2 LOF, and differential analysis revealed more accessible genomic regions in *Tet2*^-/-^ HSCs compared to HSCs isolated from WT littermate controls (Figure 1A and Table S1). Nearly half (47% of peaks) of the regions deemed to be more accessible in the *Tet2*^-/-^ HSCs were intronic (Figure 1B) and potentially related to specific gene regulation. In line with this, recent studies of chromatin accessibility revealed that HSCs with mutated *Tet2* have hypermethylation of enhancer sites^17^. Analysis of transcription factor (TF) binding sites identified specific motifs enriched in *Tet2*^-/-^ HSCs, including binding motifs for known self-renewal regulator *Smarcc1*^18^, tumor suppressor *Runx3*^19^ and cell cycle and oxidative stress regulator *Bach1*^20^, among others (Figure 1C).

**Figure 1.**
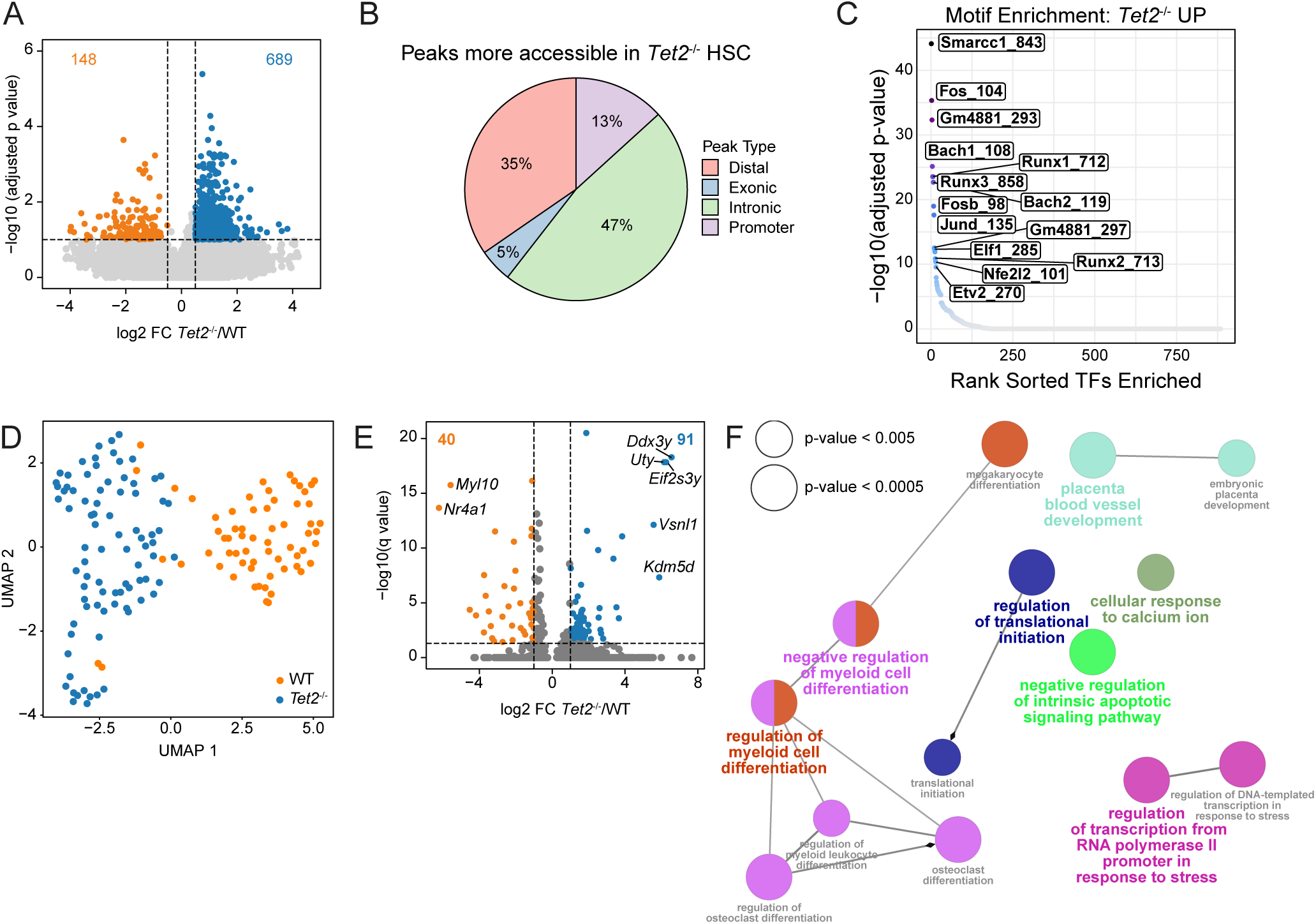
Single-cell analyses reveal increased chromatin accessibility and transcriptional activity in *Tet2*^-/-^ HSCs. (**A**) Volcano plot showing differentially accessible sites between WT and *Tet2*^-/-^ HSCs in scATAC-seq analysis (log2 FC > 0.5, adjusted p-value < 0.1). (**B**) Proportion of significantly more accessible peak regions categorized by genomic feature in*Tet2*^-/-^ HSCs. (**C**) Motif enrichment for TF motifs more accessible in *Tet2*^-/-^ HSCs. (**D**) UMAP visualization of single-cell RNA sequencing data. (**E**) Volcano plot showing differentially expressed genes between WT and *Tet2*^-/-^ HSCs in scRNA-seq analysis (log2 FC > 0.5, adjusted p-value < 0.1). (**F**) Gene Ontology Biological Processes enrichment analysis of differentially expressed genes in scRNA-seq. Node sizes reflect the statistical significance of the terms. Force directed layout presented by the Kappa score. Layout was adjusted to minimize label overlap. See also Figure S1.

Next, we undertook plate-based (scRNA-seq) to determine changes in gene expression between WT and *Tet2^-/-^* HSCs. As in our scATAC-seq data, scRNA-seq indicated significant molecular changes upon *Tet2* loss (Figure 1D and Table S2). We identified 131 differentially expressed genes (log2 fold change [FC] > 1 & adjusted p-value < 0.05), including 91 upregulated in *Tet2^-/-^* HSCs compared to WT (Figure 1E). In accordance with previous studies^21^, Gene Ontology (GO) analysis identified enrichment in pathways regulating myeloid cell differentiation and cellular response to calcium ion (Figure 1F). Together, these data suggest more active transcription in *Tet2^-/-^* ESLAM Sca-1^+^ HSCs, in agreement with a recent report^22^.

To further study the mechanisms underpinning the self-renewal differences in *Tet2^-/-^* compared to WT HSCs, scATAC-seq and scRNA-seq datasets were integrated. The closest genes to peaks identified in the scATAC-seq analysis were determined and the expression of these genes was assessed in the scRNA-seq data (distance to transcription start site [TSS] < 100,000 bp^23^). Out of 698 identified closest genes in the scATAC-seq analysis, only 18 had significantly altered expression between *Tet2^-/-^* and WT HSCs in the scRNA-seq data (Figures 2A and 2B) with 12 of these appearing amongst the top 40 differentially expressed genes in the scRNAseq-dataset. Analysis of predicted protein associations using the Genemania database^24^ showed high connectivity, strongly suggesting biological relatedness (Figure 2C). For example, among the genes with both lower chromatin accessibility and gene expression in *Tet2*^-/-^ HSCs were three members of the Krüppel-like factor (*Klf*) family (*Klf2*, *Klf4* and *Klf6*) that regulate self-renewal^25^ and *Fosb* - a member of the AP1 complex, recently shown by us to be downregulated in hibernating HSCs^26^. To further integrate information from both analyses we built a TF regulatory network using the DoRothEA database^25^. We selected all TFs whose binding motifs were significantly more accessible in *Tet2^-/-^*HSC (as shown in our scATAC-seq analysis) and curated a list of all differentially expressed genes (as determined by our scRNA-seq analysis) that the selected TFs regulate. The constructed interaction network containing all defined regulons (TF-regulatory gene pairs) identified a set of TFs potentially regulating HSC fate (Figure 2D), among which we found well-known HSC regulators such as *Gata2*, *Gata3*, *Fli1*, and *Runx1*^27–29^. Together, these data both identify factors important for normal HSC function and identify novel candidate regulators of the increased self-renewal observed in *Tet2*^-/-^ HSCs.

**Figure 2.**
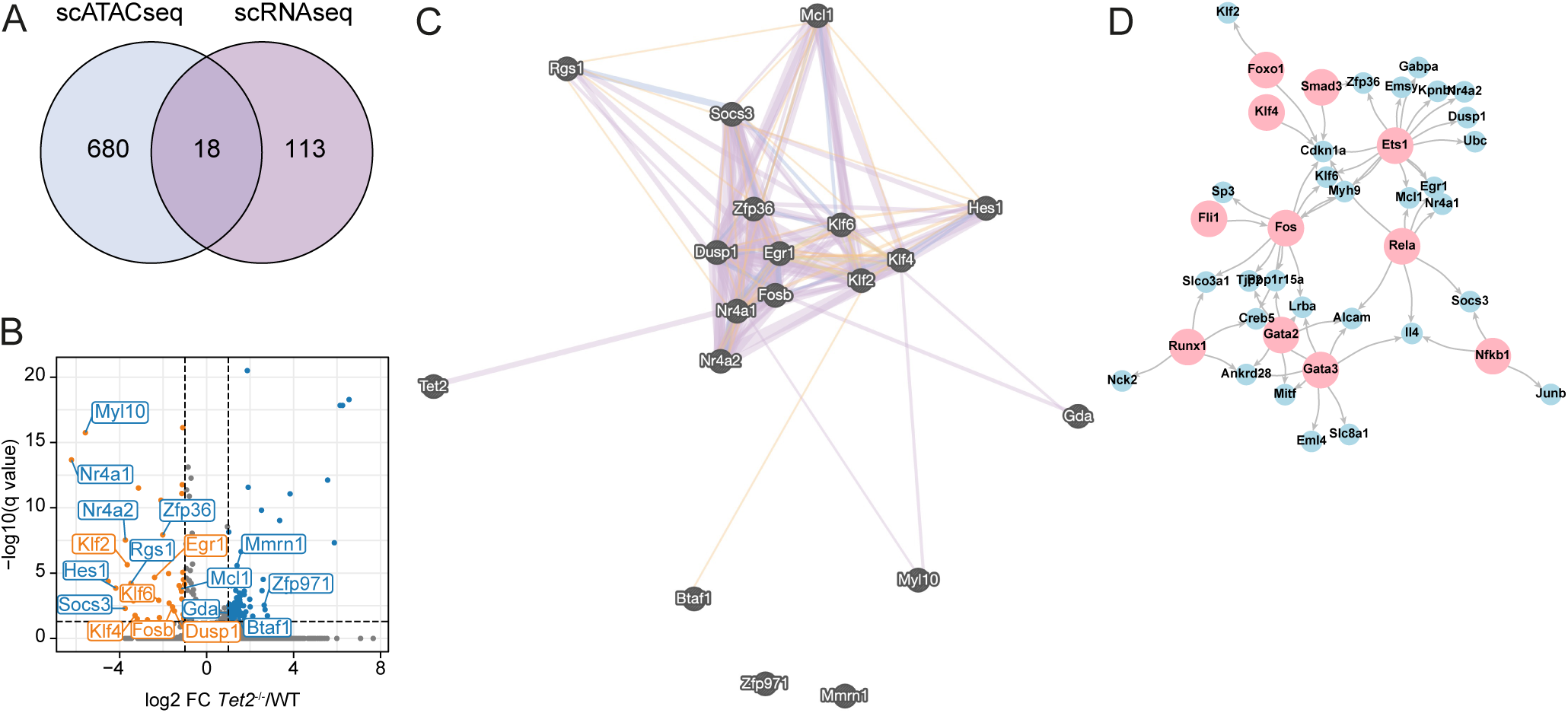
Integrative analysis of scATAC-seq and scRNA-seq datasets implicates key hematopoietic transcription factors in the regulation of *Tet2*^-/-^ HSCs. (**A**) Venn Diagram showing the overlap of targets identified in both scATAC-seq and scRNA-seq modalities (*Tet2*^-/-^ versus WT HSC pairwise testing). (**B**) Volcano plot of scRNA-seq data with labeled selected genes identified in the scATAC-seq closest gene analysis (*Tet2*^-/-^ versus WT HSC pairwise testing). Blue/orange dots indicate genes down/upregulated in scRNA-seq data analysis; blue/orange labels indicate less/more accessible markers in the scATAC-seq data analysis. (**C**) Interaction graph for targets identified in the integrative scATAC-seq/scRNA-seq data analysis constructed using the Genemania database^23^. Purple = co-expression, blue = co-localization, orange = predicted, grey = other. (**D**) Interaction graph of identified regulons - TF (scATAC-seq analysis) and genes they regulate (scRNA-seq analysis). Network built using information from the DoRothEA database^30^. Pink = TF, blue = regulated genes. See also Figure S1.

### Optimization of a low-input proteomic workflow

Our analyses of the epigenome and transcriptome of cells largely reflected current knowledge surrounding *Tet2* LOF mutations, whereby genes involved in myeloid differentiation are altered alongside a number of hematopoietic transcription factors with known roles in cellular differentiation. These data further accord with the phenotypes observed in mouse models and patients with hematological malignancies (i.e., an accumulation of myeloid progenitor cells^1,5^). In order to explore changes in populations of primary HSCs that might occur downstream of the transcriptome, a global proteomic method for low cell numbers was required. To develop this method, we used the hematopoietic progenitor cell line HoxB8-FL^31^ for protocol optimization. In an initial single-vessel sample preparation and protein-level tandem mass tag (TMT) 10plex isobaric labeling approach applied to a sample set of 3 x 10,000 cells and 7 x 20,000 cells, just 2,442 and 1,345 proteins were identified and quantified, respectively, across the entire multiplexed set (Figure 3A, 10K direct) with only modest benefit gained by increasing the smaller samples from 10,000 to 15,000 cells and carrier samples from 20,000 to 30,000 cells (Figure 3A, 15K direct), Next, through a multi-stage optimization process, we devised a method using single-vessel sample preparation, peptide-level isobaric labeling with a carrier approach, and high-resolution offline fractionation which resulted in robust quantification of over 3,500 proteins from as few as 10,000 cells (Figures 3A-3C).

**Figure 3.**
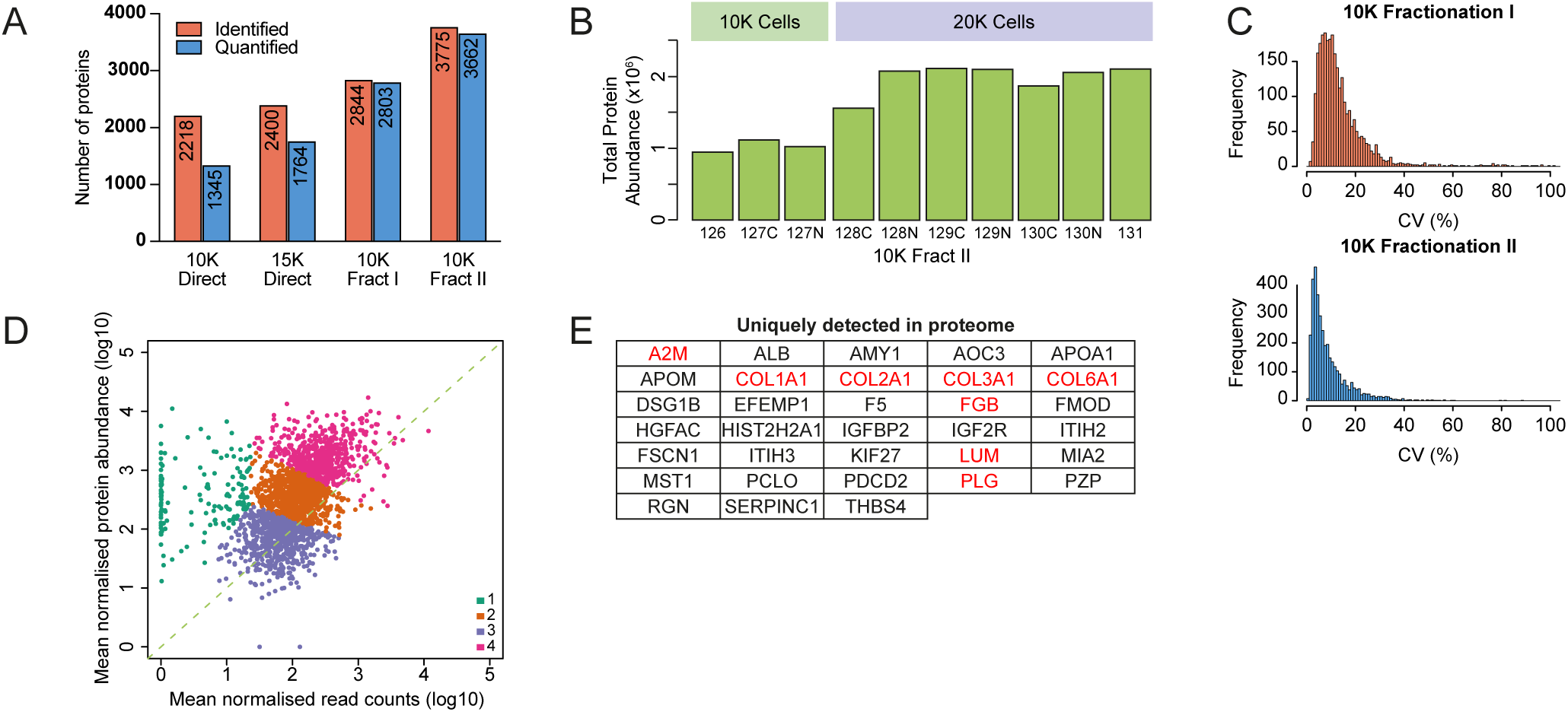
An optimized workflow enables deep proteomic profiling of low numbers of hematopoietic cells. (**A**) Number of identified and quantified proteins in HoxB8-FL cells using direct injection or offline fractionation into 6 fractions (Fract I) or 5 fractions × 2 runs each with different upper intensity precursor selection limit in the two runs (Fract II). (**B**) Total protein abundance across TMT channels in the 10K Fract II experiment from (**A**). (**C**) Coefficient of variance (CV) values for the 10K Fract I and 10K Fract II experiments. (**D**) Correlation between protein and mRNA abundance in HoxB8-FL cells. Points are colored according to the k means cluster they belong to. (**E**) Identifications unique to proteome data. Proteins in red are associated with ECM organization.

To assess the type of information gained by proteome-level characterization, we mapped protein abundance from the Fract II experiment (Table S3) against gene expression^32^ (Figure 3D) and observed a positive correlation overall. Clustering analysis identified a set of genes/proteins showing particularly strong enrichment at the protein level (Figure 3D, cluster 1 shown in green), including 33 that were uniquely detected in the proteome data (Figure 3E). Intriguingly, 8 of these 33 proteins (24%) were associated with ECM organization (Figure 3E, highlighted in red), suggesting that ECM components might be more readily captured by proteomic relative to transcriptomic analysis.

### Global proteomics identifies distinct molecular changes in *Tet2^-/-^* HSPCs not captured by transcriptomics

To generate a comprehensive molecular map of WT and *Tet2*^-/-^ HSPCs, we applied our low-input proteomic workflow (Figure 3) to WT and *Tet2*^-/-^ HSPCs with durable (Lineage [Lin]^-^ cKit^+^ CD45^+^ CD48^-^ CD150^+^; collectively called “CD150^+^”) or finite (Lin^-^ cKit^+^ CD45^+^ CD48^-^ CD150^-^; collectively called “CD150^-^”) self-renewal^33^ (Figure 4A). We additionally analyzed the proteome of WT ESLAM HSCs (CD45^+^ CD48^-^ CD150^+^ EPCR^+^) as a reference LT-HSC population^33^ and WT Lin^-^ cKit^+^ (LK) cells as a carrier proteome population to increase mapping efficiency^9^. To assess the degree of post-transcriptional regulation and the overall correlation between transcriptome and proteome in *Tet2*^-/-^ HSPCs, we additionally performed bulk RNA-seq on WT and *Tet2*^-/-^ CD150^+^ and CD150^-^ cells. Using just 10,000-30,000 cells per sample, the MS analysis identified 4,133 unique proteins, out of which 3,989 (∼97%) were reliably quantified across all cell populations (Table S4). Notably, our low cell number multiplex captured 55% of proteins previously quantified in HSCs^34^ using only 2.5-7.5% of total cell input per sample. Normalization against sample loading successfully corrected for the difference in cell number between samples (Figure S2A).

**Figure 4.**
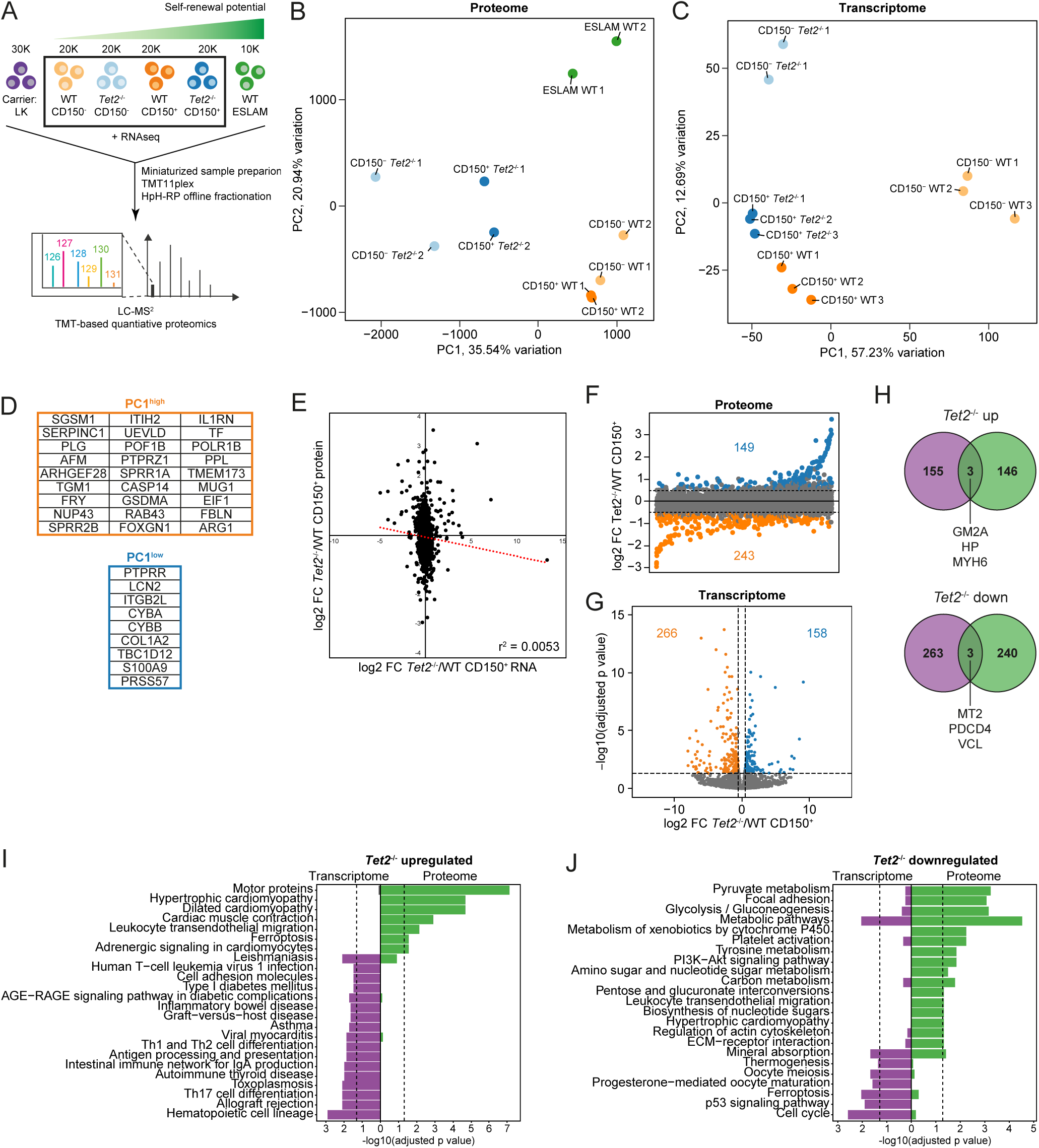
The proteome and transcriptome highlight alterations in disparate cellular processes upon loss of *Tet2* in HSPCs. (**A**) Workflow for LC-MS/MS analysis of WT and *Tet2*^-/-^ HSPCs. (**B, C**) PCA of proteomic (**B**) and transcriptomic (**C**) data. (**D**) Top 15% loadings from PC1 in (**B**). (**E**) Correlation between protein and gene expression differences (log2 FC) between *Tet2*^-/-^ and WT CD150^+^ cells. The dotted red line indicates the linear trendline. *r^2^* represents the Pearson correlation coefficient. (**F**) Protein expression difference between *Tet2*^-/-^ and WT CD150^+^ cells. Candidate target proteins (log2 FC > 0.5 across all comparisons between *Tet2*^-/-^ and WT CD150^+^ cells) enriched and depleted in *Tet2*^-/-^ relative to WT CD150^+^ cells are shown in blue and orange, respectively. (**G**) Gene expression difference between *Tet2*^-/-^ and WT CD150^+^ cells. Candidate target genes (log2 FC > 0.5 and adjusted p-value < 0.05) enriched and depleted in *Tet2*^-/-^ relative to WT CD150^+^ cells are shown in blue and orange, respectively. (**H**) Overlap between candidate target genes and proteins enriched or depleted in *Tet2*^-/-^ relative to WT CD150^+^ cells. (I, J) KEGG pathway analysis of candidate target proteins and genes enriched (**I**) or depleted (**J**) in *Tet2*^-/-^ relative to WT CD150^+^ cells. The dotted lines mark adjusted p-value = 0.05 See also figure S2.

Following the generation of global proteomic datasets for WT and *Tet2*^-/-^ HSPCs, we first compared proteome and transcriptome datasets. Principal component analysis (PCA) of proteomic data clearly separated cell populations according to genetic background in principal component (PC) 1 (Figure 4B), whereas separation in PC1 and PC2 for RNA-seq data was driven by cell type rather than *Tet2* mutational status (Figure 4C), indicating that the transcriptome and proteome have substantial global differences in regulation. Interestingly, in Figure 4B, the *Tet2^-/-^* CD150*^-^* samples are most distinct compared to WT populations and *Tet2^-/-^* CD150*^+^* cells, suggesting that proteome dysregulation in *Tet2^-/-^* hematopoietic cells is exacerbated as cells mature past the stem cell state, a finding potentially associated with the myeloid skewing observed with loss of TET2^1,5^. To a lesser extent, this was also reflected in PC2 of the transcriptome data (Figure 4C). To investigate which proteins best separated WT and *Tet2^-/-^* cells, we extracted the top and bottom 15% of loadings for PC1 (27 WT-enriched and 9 *Tet2^-/-^*-enriched; Figure 4D). Proteins associated with the TET2-deficient cell populations included inflammatory proteins S100A9 and LCN2, which have been reported to be involved in the pathogenesis of myelodysplastic syndrome (MDS) and myelofibrosis (MF), respectively^35,36^. WT-enriched loadings included interleukin-1 (IL-1) receptor antagonist protein IL-1RA, a protein important for dampening IL-1-driven inflammation, which was recently shown to contribute to the clonal outgrowth of *Tet2^+/-^* HSPCs during aging^37^. Two of the top loadings for WT cells in PC1 were anti-coagulant proteins antithrombin III (SERPINC1) and plasminogen (PLG), suggesting potential dysregulation of clotting mechanisms in *Tet2^-/-^* HSPCs. Intriguingly, LOF mutations in *TET2* have recently been linked to an increased risk for MPN-associated thrombosis, and patients carrying *TET2* mutations have significantly lower levels of antithrombin III than those with intact *TET2* expression^38^. Our data suggest that the increased production of inflammatory proteins and impaired clotting functions associated with mutations in *Tet2* are evident already at the level of HSCs.

Next, we specifically interrogated the immature CD150^+^ population. There was no correlation between the protein and RNA datasets for the CD150^+^ population (*r*^2^ = 0.0053; Figure 4E) which accords with previous studies that reported a high degree of post-transcriptional regulation in early HSPCs^10,13,14^, and poor correlation between proteome and transcriptome in HSCs compared to downstream progenitors^13^. The correlation between transcriptome and proteome in the CD150^-^ population was equally poor (*r*^2^ = 0.0056; Figure S2B). To further assess concordance between the two datasets, we next generated shortlists of candidate proteins and genes and compared the lists (Figures 4F and 4G). In the CD150^+^ LT-HSC-enriched population, only 6 candidate targets overlapped between the two datasets; these included inflammatory biomarker haptoglobin (HP) which showed elevated expression in *Tet2*^-/-^ cells, and anti-inflammatory methallothionein (MT) 2, which was enriched in WT relative to knockout cells (Figure 4H). The overlap in the CD150^-^ population was substantially higher with 33 and 97 candidate proteins (representing 15 and 21% of all candidate proteins, respectively) significantly enriched at the transcript level in *Tet2*^-/-^ and WT cells, respectively (Figure S2C).

Despite the low overlap of specific targets between the MS and RNA-seq datasets, KEGG pathway enrichment analysis of candidate proteins/genes in the CD150^+^ LT-HSC-enriched population highlighted some commonalities (Figures 4I and 4J), suggesting significant dysregulation in HSPCs upon loss of *Tet2*. The most prominent of these categories were “metabolic pathways” and “mineral absorption”, which were both strongly enriched among proteins and genes downregulated in *Tet2*^-/-^ relative to WT CD150^+^ cells. Surprisingly, the top enriched pathway among proteins upregulated in *Tet2*^-/-^ relative to WT cells was “motor proteins”, and the proteomic datasets highlighted changes in additional pathways related to the actomyosin motor and the ECM upon loss of TET2 in immature HSPCs, such as “leukocyte transendothelial migration”, and “ECM-receptor interaction” (Figure 4J). Metabolism and the actomyosin motor were also altered in CD150^-^ cells, pointing to their dysregulation as a general feature of TET2 loss in HSPCs (Figures S2D and S2E).

### ECM interactions regulate self-renewal of *Tet2*^-/-^ HSCs

Reactome pathway enrichment analysis (Figures 5A and 5B) identified several pathways related to the actomyosin motor and to ECM organization among proteins more highly expressed in either *Tet2*^-/-^ (Figure 5A) or WT (Figure 5B) cells. Analysis of individual proteins contained within the “Extracellular matrix organization” pathway revealed distinct expression patterns of ECM proteins in the three most HSC-enriched cell populations in our proteomic data (Figures 5C), with proteins either being depleted or enriched with increasing self-renewal potential. A subset of these proteins, including four collagen family members, displayed particularly high expression in *Tet2*^-/-^ CD150^+^ cells, and three of these (ITGB2, COL2A2, and P4HB) overlapped with the more highly HSC-enriched WT ESLAM population. Notably, bulk RNA-seq data did not capture these changes, instead showing that ECM-associated genes were either unchanged between WT and TET2-deficient cells or had opposing expression patterns compared to the proteome data (Figure 5D).

**Figure 5.**
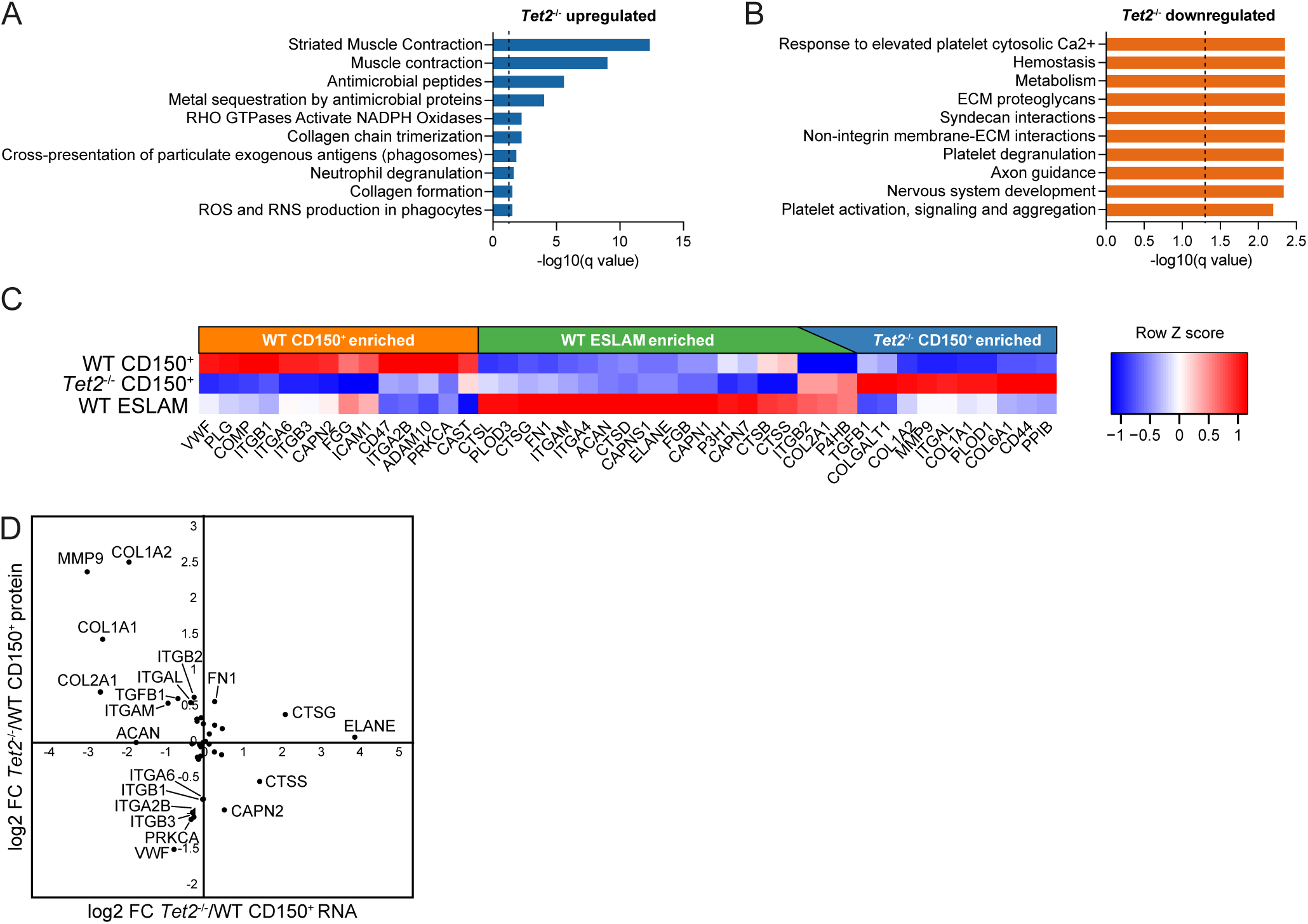
Expression of ECM proteins is altered upon loss of *Tet2* and correlates with self-renewal potential. (**A, B**) Reactome pathway analysis of candidate target proteins enriched (**A**) or depleted (**B**) in *Tet2*^-/-^ relative to WT CD150^+^ cells. The dotted lines mark q-value = 0.05. (**C**) Relative abundance of proteins contained within the Reactome pathway “Extracellular matrix organization” in WT CD150^+^, *Tet2*^-/-^ CD150^+^, and WT ESLAM cells. (**D**) Correlation between protein and gene expression differences (log2 FC) of proteins/genes contained within the Reactome pathway “Extracellular matrix organization” between *Tet2*^-/-^ and WT CD150^+^ cells.

All ECM proteins identified in the MS data are known interaction partners, and, intriguingly, proteins enriched in *Tet2*^-/-^ and WT CD150^+^ cells were positioned in distinct areas of the interaction network (Figure 6A). We next used flow cytometry to assess surface expression of integrins CD44, ITGB1/CD29, ITGB3/CD61, and ITGA2B/CD41, which were all among proteins differentially abundant between *Tet2*^-/-^ and WT CD150^+^ cells (Figures 6A-6E). Intriguingly, *Tet2^-/-^* ESLAM HSCs showed increased surface expression of all four proteins relative to WT cells (Figures 6B-E). The disparity between cellular proteome data and surface expression of ITGB1/CD29, ITGB3/CD61 and ITGA2B/CD41 may indicate differential localization of these proteins in healthy and pre-leukemic HSCs. Critically, ITGA2B/CD41, a marker for myeloid-biased HSCs^39^, was nearly undetectable on the surface of WT HSCs while expression could be detected in all TET2-deficient samples (Figure 6E). Integrin receptors are heterodimeric, and one such heterodimer is formed by ITGA2B/CD41 and ITGB3/CD61. To assess the degree of co-localization of ITGA2B/CD41 and ITGB3/CD61, we employed state-of-the-art imaging FACS^40^ coupled with a downstream functional assay of *Tet2*^-/-^ and WT ESLAM HSCs (Figures 6F-6H). We observed a significantly higher correlation between ITGA2B/CD41 and ITGB3/CD61 in *Tet2*^-/-^ and WT HSCs (Figure 6G), suggesting increased co-localization of these two integrins in pre-leukemic HSCs. Critically, the correlation was highest in the *Tet2*^-/-^ HSCs that gave rise to small clones in culture, which have been previously shown to be highly enriched for HSCs^41^ (Figure 6H).

**Figure 6.**
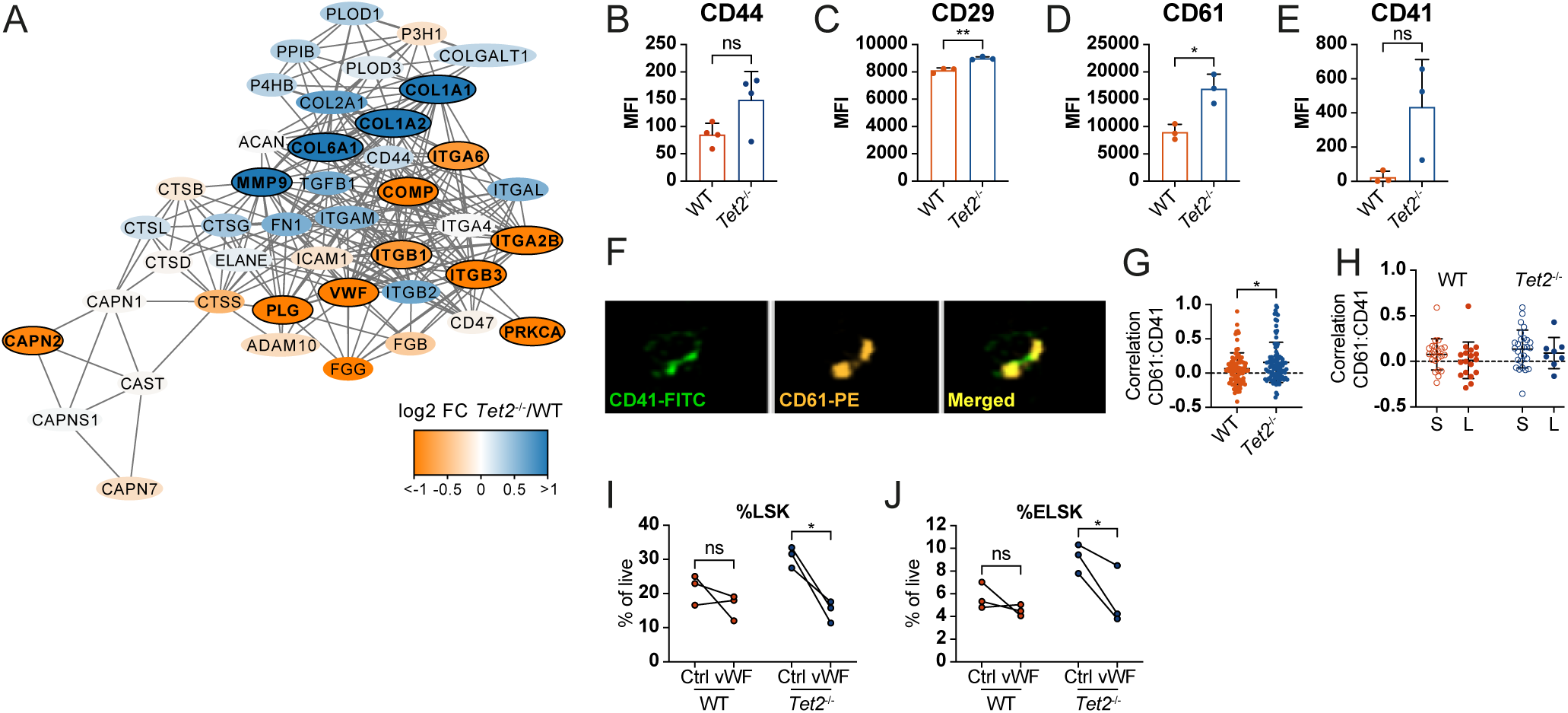
Interaction with niche-anchored vWF and HA inhibits expansion of *Tet2*^-/-^ HSCs. (**A**) Protein interaction network of proteins contained within the Reactome pathway “Extracellular matrix organization”. Outlined nodes represent candidate target proteins. (**B**-**E**) Surface expression of CD44 (**B**), CD29 (**C**), CD61 (**D**), and CD41 (**E**) on WT and *Tet2*^-/-^ HSCs. MFI = median fluorescent intensity. (**F**) Representative images of CD41 and CD61 expression in *Tet2*^-/-^ HSCs. (**G**) Correlation between CD61 and CD41 in index-sorted WT and *Tet2*^-/-^ HSCs. (**H**) Correlation between CD61 and CD41 in small (S; 50-5,000 cells) and large (L; >10,000 cells)^41^ WT and *Tet2*^-/-^ HSC clones following culture for 11 days. (**I**, **J**) Frequency of HSPCs (LSK; **I**) and HSCs (ELSK^42^; **J**) cells following 28-day culture of WT and *Tet2*^-/-^ ESLAM HSCs on hydrogels functionalized with vWF. **p < 0.01, *p < 0.05 and ns= non-significant. Error bars = SD. See also Figure S3.

The differential expression of ECM proteins and their integrin receptors, both in the cellular proteome and on the cell surface, led us to hypothesize that *Tet2^-/-^* and WT HSCs may differ in their ability to interact with the niche, which in turn could affect important cellular functions, including survival, self-renewal capacity, and differentiation. In order to test the functional impact of different ECM components, we cultured ESLAM HSCs derived from WT and *Tet2*^-/-^ animals on STEMBOND hydrogels functionalized with vWF, which is depleted from *Tet2^-/-^* relative to WT HSCs or hyaluronan (HA) as HA receptors CD44 and ITGB1/CD29 were among proteins showing self-renewal and *Tet2*-status-associated differences in expression (Figures 5C and 6A-6C). As expected, loss of TET2 confers a self-renewal advantage in HSCs^1,5^ that presents itself as an increase in the frequency of Lin^-^Sca-1^+^cKit^+^ (LSK) HSPCs and EPCR^+^ LSK (ELSK) HSCs^42^ compared to WT controls (Figures 6I and 6J, Figures S3A and S3B). Strikingly, this mutant self-renewal advantage was abolished when cells were cultured in the presence of hydrogel-anchored vWF (Figures 6I and 6J), and, to a lesser extent, HA (Figures S3A and S3B). The reduced fraction of progenitors (LSK) and HSCs (ELSK) was only observed in the *Tet2^-/-^* cultures, suggesting that vWF is selectively (and negatively) influencing TET2-mediated HSC expansion (Figures 6I and 6J). Together these data show that exposure to different ECM can regulate self-renewal capacity of *Tet2*^-/-^ HSCs and firmly establish that features captured exclusively by proteomic analysis provide novel insight into HSC biology.

## Discussion

TET2 has been widely studied in the context of clonal hematopoiesis and myeloid malignancies due to its recurring LOF mutations and its functional role in HSC self-renewal. The exact molecular mechanism of the increased self-renewal initiated by loss of TET2 function remains unclear and transcriptomic studies to date have been unable to identify clear drivers of increased HSC self-renewal. Our integrated ATAC-seq and scRNA-seq analyses implicated KLF family members and key hematopoietic transcription factors *Gata2*, *Gata3*, *Fli1*, and *Runx1* in the regulation of TET2-mutant HSCs. Then, moving beyond transcriptomics to capture global proteome level data, we identify a distinct set of ECM molecules with specific roles in altering the function of *Tet2*-mutated HSCs. Intriguingly, in HSCs as well as in hematopoietic cell line HoxB8-FL, ECM protein abundance does not correlate with expression of associated transcripts as determined by RNA-seq, suggesting a high degree of post-transcriptional regulation in this group of proteins.

Functionalizable hydrogels represent a novel tool with which to study components of the ECM and their impact on cell function. In this study, we utilize STEMBOND hydrogels^43^, which permit robust matrix tethering and have tunable stiffness, to test the ECM components that emerged from our proteomic studies. This permits investigation of biophysical properties of cells with LOF *Tet2* mutations and we demonstrate a clear role for vWF in specifically restricting TET2-mediated HSC expansion *in vitro*. Furthermore, our proteomic data point to actomyosin motor control as being dysregulated when *Tet2* is mutated. While TET2 has been previously implicated in cytoskeleton organization in ovarian cells^44^ and in smooth muscle cell plasticity^45^, its role in the actomyosin motor of HSPCs has not yet been described. In this vein, myosins are upregulated during inflammatory stress in HSCs^14^, and their inhibition impairs growth and survival of acute myeloid leukemia (AML) cells^46^, implying that their high expression in *Tet2* mutant HSPCs may be linked to the aberrant phenotype of the cells.

We also found that TET2-deficient HSCs have altered expression of several integrins, both in the cellular proteome and on the cell surface. Both ITGA2B/CD41 and ITGB3/CD61 have been shown to mark myeloid-biased HSCs^39^, and CD61 was recently shown to identify HSCs with enhanced self-renewal potential during aging^47^. Notably, antibody-based depletion of myeloid-biased HSCs has recently emerged as a strategy for rejuvenation of the hematopoietic system^39^. Our data indicate that a similar approach may selectively deplete *Tet2*-mutant HSCs while sparing healthy cells and could represent a novel therapeutic strategy for purging pre-malignant HSCs from the bone marrow. CD61 is one of the receptors for vWF^48^ and thus the increased surface expression on *Tet2*-mutant cells could also explain the sensitivity of these cells to vWF, which is depleted in TET2-deficient relative to WT HSCs. These results highlight that mutant HSC self-renewal is regulated by an intricate balance between surface level expression of receptors and abundance of corresponding ECM proteins.

There were also a number of proteins more highly expressed in WT cells, in particular a set of proteins related to platelet function. Interestingly, the WT-enriched proteins representing these processes include those with anti-coagulant (e.g. PLG, SERPINC1 and ANXA5) as well as pro-thrombotic (e.g. FGG, VWF, and ITGB3) functions, suggesting that TET2 loss results in a general decrease in expression of proteins related to platelet production and coagulation in HSPCs and perhaps even related to a shift in HSC subtypes away from megakaryocyte-biased HSCs.

Overall, our study emphasizes the importance of moving beyond transcriptomic studies to reveal new aspects of mutant cell biology during processes of HSC self-renewal and leukemogenesis. In particular, proteomic studies have triggered the investigation of the mechanisms by which the ECM alters HSC self-renewal and influences clonal advantage competition during aging and disease. How changes in ECM composition throughout ageing might contribute to the clinical observations of clonal hematopoiesis and pre-leukemic cell expansion is an intriguing concept that accords with recent studies showing that integrins and their molecular regulators underpin healthy ageing^47,49,50^. This in turn opens up new lines of thinking regarding potential therapies and new tools such as functionalizable hydrogels will accelerate discoveries that reach well beyond the HSC system for applications in numerous other stem cell systems as has already been pioneered for oligodendrocyte precursors^51^ and pluripotent stem cells^43^.

## Acknowledgments

Work in the Prof. DG Kent laboratory was supported by a European Research Council Starting Grant (ERC-2016-STG-715371), a Cancer Research UK Programme Foundation Award (DCRPGF\100008), an MRC-AMED joint award (MR/V005502/1), the UK Medical Research Council (MC_PC_21043; MR/Y011945/1), and Blood Cancer UK (24014). The Kent lab is also supported by the National Institute for Care and Health Research Leeds Biomedical Research Centre (NIHR203331). Dr. M Jassinskaja was supported by a Swedish Research Council International Postdoc grant (2021-00185), and project grants from The Royal Physiographic Society of Lund and Siv-Inger och Per-Erik Anderssons minnesfond. M Gonka was the recipient of a Biotechnology and Biological Sciences Research Council White Rose DTP PhD Studentship (BB/T007222/1). Dr. MS Shepherd was the recipient of a Biotechnology and Biological Sciences Research Council Industrial Collaborative Award in Science and Engineering (iCase) PhD Studentship. Dr. JLC Che was supported by an MRC PhD Studentship under the University of Cambridge Doctoral Training Programme. Dr D Bode was supported by a Wellcome PhD Studentship. Work in Cambridge was further supported by core support grants by the Wellcome and Medical Research Council (MRC) to the Wellcome-MRC Cambridge Stem Cell Institute (203151/Z/16/Z). Work in the Prof. B Göttgens laboratory was supported by Wellcome (206328/Z/17/Z and 203151/Z/16/Z), Blood Cancer UK (18002), Cancer Research UK (C1163/A21762), and UKRI Medical Research Council (MC_PC_17230). The authors would like to thank Professor Anjana Rao for originally providing the Tet2^-/-^ mouse, and Drs Veronique Voisin, Andrew Zeng, Matthew Care and Alastair Droop for their valuable input on scRNA-seq and scATAC-seq analyses. The authors also greatly acknowledge the expert technical assistance provided by the Genomics, Imaging & Cytometry, and Metabolomics and Proteomics Facilities within the Biosciences Technology Facility at the University of York, and the CIMR Flow Cytometry core (Reiner Schulte, Chiara Cossetti, and Gabriela Grondys-Kotarba). We also thank Tina Hamilton and Dean Pask for technical assistance and the staff of the animal facilities at the University of York and University of Cambridge. The Viking cluster was used during this project, which is a high performance computing facility provided by the University of York. We are grateful for additional computational support from the University of York, IT Services and the Research IT team.

## Author contributions

DGK, JC, and BG designed the study together with DB and MJ. DB and IK performed scRNA-seq and scATAC-seq experiments under supervision of NKW and BG. DB and TIR performed proteome analysis under supervision by JC. DB and MG analyzed scRNA-seq data with SM. MG analyzed scATAC-seq data with BT, MSV, ANH and SA. MG performed integrative analysis of scRNA-seq and scATAC-seq data with BT and BAH. MJ and DB analyzed proteome data. MB, MJ, JLCC, MSS, AJH, EB, JM, MG, JARL and LCC performed HSC isolation, flow cytometry and cell culture experiments with experimental design input from SY and WJB. MJ analyzed all flow cytometry data. GV, HK, ACH, EB and JM performed animal work. MJ prepared the figures. DGK supervised the study. MJ, MG, and DGK wrote the paper together with DB, AJH, EB, JM and input from other authors.

## Supplemental information titles and legends

Figure S1. Related to Figures 1 and 2.

Figure S2. Related to Figure 4.

Figure S3. Related to Figure 6.

Table S1. scATAC-seq data of WT and *Tet2*^-/-^ HSCs, related to Figure 1.

Table S2. scRNA-seq data of WT and *Tet2*^-/-^ HSCs, related to Figure 1.

Table S3. Proteome of HoxB8-FL cells, related to Figure 3.

Table S4. Proteome and bulk transcriptome of *Tet2*^-/-^ and WT HSCs, related to Figure 4.

## Figure legends

**Figure S1. Quality control of scATAC-seq and scRNA-seq data. Related to Figure 1** (A, B) Fragment size distribution (A) and fragment distribution across samples (B) in scATAC-seq data. (C) Motif enrichment for TF motifs more accessible in WT HSC in scATAC-seq. (D) Number of genes detected in each cell (nFeature_RNA), total number of molecules detected within a cell (nCount_RNA), and fraction of reads mapped to mitochondrial genes (percent.mt) in scRNAs-eq data. (E) Elbow plot for scRNAseq data. The top 5 PCs were selected for the downstream analysis.

**Figure S2. Proteome of CD150^-^ short term (ST)-HSCs undergoes extensive remodeling upon loss of TET2. Related to Figure 4.** (A) Normalized intensity across *Tet2*^-/-^ and WT HSPC samples analyzed by LC-MS/MS. (B) Correlation between protein and gene expression differences (log2 FC) between *Tet2*^-/-^ and WT CD150^-^ cells. The dotted red line indicates the linear trendline. *r^2^* represents the Pearson correlation coefficient. (C) Overlap between candidate target genes and proteins enriched or depleted in *Tet2*^-/-^ relative to WT CD150^-^ cells. (D, E) KEGG pathway analysis of candidate target proteins and genes enriched (D) or depleted (E) in *Tet2*^-/-^ relative to WT CD150^-^ cells. The dotted lines mark adjusted p-value = 0.05.

**Figure S3. Impact of hyaluronan on expansion of *Tet2*^-/-^ HSCs in culture. Related to Figure 6.** (A, B) Frequency of HSPCs (LSK; A) and HSCs (ELSK; B) following 28-day culture of WT and *Tet2*^-/-^ HSCs on hydrogels functionalized with hyaluronan (HA). ns = non-significant.

## Methods

### Mice

Wild-type C57BL/6N and *Tet2*^-/-^ mice bred in-house were used for all experiments. Adult animals aged 12-16 weeks were used for all experiments except for MS analysis where used animals were 30 weeks old. Animals were housed in individually ventilated cages (IVC) and provided with sterile food and water ad libitum. All mice were kept in specified pathogen-free conditions, and all procedures performed according to the United Kingdom Home Office regulations, in accordance with the Animal Scientific Procedure Act.

### Flow cytometry and FACS

For sorting of primary HSPCs, bone marrow was extracted from hind limbs, hips, sternum, and spine collected in ice cold phosphate buffered saline (PBS). Bones were crushed using a mortar and pestle and cell suspension was mechanically dissociated using a pipette and passed through a 40 µM filter. Red blood cells were lysed by incubation with ammonium chloride (Stem Cell Technologies). Mature cells were magnetically depleted from the cell suspension using the EasySep Mouse Hematopoietic Stem and Progenitor Cell Isolation Kit (Stem Cell Technologies). For sorting for MS analysis, cell suspensions were stained with fluorophore-conjugated antibodies against CD45, CD150, CD48, EPCR, and cKit by incubation for 30 min on ice protected from light. For all other experiments, anti-cKit antibody was omitted and anti-Sca-1 antibody was included. In all flow cytometry and FACS experiments, 7-aminoactinomycin D (7-AAD) staining was used to exclude dead cells. FACS experiments were performed on a BD Influx at the Cambridge Institute for Medical Research, or a Beckamn Coulter MoFlo Astrios or BD FACS Discoverer S8 at the Imaging & Cytometry Technology Facility at the University of York. For flow cytometric analysis of cultured HSCs, cell suspensions were stained with fluorophore-conjugated antibodies against CD45, CD11b, Gr-1, cKit, Sca-1, and EPCR. Flow cytometric analysis was performed on a Beckman Coulter CytoFlex LX or BD LSRFortessa X20 at the Imaging & Cytometry Technology Facility at the University of York.

### HSC gel culture

96-well CELLview™ plates (Greiner) were activated to allow the binding of StemBond. Plates were treated inside a plasma system (Henniker HPT-200) and functionalized using 5% Bind Silane solution (GE Healthcare). Plates were washed thoroughly with 100% ethanol. 3 ml soft hydrogel solutions were prepared using 40% acrylamide (210 µl), 2% Bis-acrylamide (120 µl), TEMED (15 µl), 10% Ammonium Persulfate (APS, 30uL), and water (2461.8 µl) and transferred to the CELLview™ plates. Following polymerisation, gels were rinsed twice in methanol, followed by a PBS rinse. Prior to activation with EDAC/NHS solution (Sigma Aldrich), gels were rinsed with pH 6.1 MES buffer. Once activated, gels were rinsed with chilled 60% methanol in PBS, followed by a 50 mM pH 8.5 HEPES buffer rinse. Gels were coated 100-200 µg/ml of ECM protein diluted in HEPES buffer and incubated overnight at 4℃. Following incubation, the protein solution was removed and gels were rinsed with HEPES buffer. Ethanolamine solution (0.5 M; ChemCruz) in HEPES buffer was used to block the gels for 30 minutes at room temperature. Gels were rinsed for a final time with pH 7.4 HEPES buffer and PBS to equilibrate the pH. Gels were stored at 4℃ until use. 50 ESLAM HSCs per well were sorted directly onto gels into HSC expansion media^53,54^ and maintained in culture at 37 C and 5% CO2 for 28 days with media changes every 2-3 days.

### Bulk RNA-seq

To match the cell populations extracted for proteomic profiling, Lin^-^ CD45^+^ CD48^-^ CD150^+^ cKit^+^ (collectively called CD150^+^) and Lin^-^ CD45^+^ CD48^-^ CD150^-^ cKit^+^ (collectively called CD150^-^) were isolated using FACS (as described above). A total of 1000 cells were collected per biological sample from homogenized cell extracts (femurs, tibiae, hips and spines) of *Tet2*^-/-^ and WT mice. RNA extraction was performed using the Picopure RNA Isolation Kit (Thermo Scientific) according to manufacturer’s protocol. Library preparation and sequencing was performed at the Cancer Research UK Cambridge Institute Genomics Core as previously described^42^. Data processing was conducted as previously described^42^. In brief, adapter trimming was performed using trim_galore (parameters: --paired --quality 30 --clip_R2 3). Reads alignment against the Mus musculus genome build (mm10) was conducted using STAR (default parameters). Gene counts were computed using HTSeq (parameters: --format = bam – stranded = reverse –type = exon –mode = intersection-nonempty --additional-attr = gene_name). Downstream processing and quality control was conducted using EdgeR (version 3.28.1,https://pmc.ncbi.nlm.nih.gov/articles/PMC9535767/#embr202255502-bib-0118, https://pmc.ncbi.nlm.nih.gov/articles/PMC9535767/#embr202255502-bib-0117), with read counts being transformed to counts per million (cpm), genes with fewer than 2 samples expressing >1cpm being excluded and read count normalization being performed using the trimmed mean of M values (TMM) method^55^.

### scATAC-seq

scATAC-seq data has been generated from *Tet2*^-/-^ mice and WT littermate controls. Bone marrow was lineage depleted and CD48^-^CD150^+^CD45^+^EPCR^+^ (ESLAM) HSCs were sorted. Cells were isolated from 5 WT mice and 5 *Tet2*^-/-^ mice (4,000 cells per genotype). Libraries were prepared using the 10x Genomics Chromium Next GEM Single Cell ATAC Reagent Kits v1.1. Sequencing was run at Leeds University Next Generation Sequencing facility using a NextSeq 2000. Read alignment to a reference genome (mm10) was performed using CellRanger pipeline (CellRanger-ATAC, 10x Genomics, cellranger-atac count). For downstream data analyzes, ArchR workflow was used^56^. Due to the low intra-sample heterogeneity, the ArchR’s simulation of synthesized *in silico* doublets over the data was not used to exclude potential doublet cells and doublets were removed by filtering cells containing less than 40,000 fragments instead. The term frequency-inverse document frequency (TF-IDF) normalization and the singular value decomposition (SVD) were performed (latent semantic indexing, LSI^57^) using ArchR’s addIterativeLSI. Uniform Manifold Approximation and Projection (UMAP) dimension reduction^58^ was run with ArchR’s addUMAP. Pseudo-bulk replicates were created, and peaks were called using MACS2^59^. Marker peaks unique to individual groups were identified with ArchR’s getMarkerFeatures. Paired samples Wilcoxon test was used to compare WT and *Tet2*^-/-^ samples (FDR 0.1 & absolute log2 FC > 0.5)). Transcription factor binding motifs were annotated using ArchR’s addMotifAnnotations function, and motif set from the cisbp database^60^ was used (chromVAR package^61^). Differentially accessible peaks were tested for motif enrichment with ArchR’s peakAnnoEnrichment function. Closest genes to the accessible regions were identified if the distance to the transcription start site was < 100k base pairs (FDR <= 0.1 & log2 FC >= 0.5). Computational analysis was performed using the University of York Research High Performance Computing Cluster (Rocky 8.8, Viking2). Plots were made with ArchR^56^, ggplot2^62^, Seurat^63,64^, Cytoscape^65^ and ClueGO^66^.

### Plate-based scRNA-seq

scRNA-seq data were generated from *Tet2*^-/-^ mice and WT littermate controls. ESLAM HSCs were FACS-purified as previously described. Freshly isolated HSCs were subjected to single-cell RNA SmartSeq2 sequencing (2 x 96-well plates, WT and *Tet2*^-/-^ mice, one mouse per plate). RNA was extracted using the Picopure RNA isolation kit (Thermo Fisher). Libraries were prepared using a protocol adapted from the msSCRB-seq workflow and quality control was performed using the Bioanalyzer system (Agilent). ERCC external RNA Spike-In controls were used (ERCC RNA Spike-In Mix; ThermoFisher). Constructed libraries were sequenced using the Novogene system using a paired end 150 bp run on a NovaSeq S4 flowcell. Sequenced reads were aligned to the mm10 reference mouse genome using STAR aligner^67^ and gene counts were computed using featureCounts^68^. For downstream scRNA-seq data analysis, the Seurat workflow^63,64^ was used. Data were normalized using regularized negative binomial regression (Seurat’s SCTransform). Principal component analysis (PCA) reduction analysis was performed with Seurat’s RunPCA (default parameters), and the top 5 principal components were selected. UMAP^58^ was run with Seurat’s RunUMAP (default parameters). Local neighbourhoods were defined with Seurat’s FindNeighbors, and cells were clustered using the Louvain algorithm^69^ (Seurat’s FindClusters). Differentially expressed genes for *Tet2*^-/-^ and WT cell groups were found with Seurat’s FindMarkers. Significantly up/down regulated genes were defined if Q-value < 0.05 & absolute average log2 FC > 1. To compare functional profiles for identified genes, clusterProfiler^70^ and enrichplot^71^ were used. Computational analysis was performed using the University of York Research High Performance Computing Cluster (Rocky 8.8 and Viking2). Plots were made with ArchR^56^, ggplot2^62^, Seurat^63,64^, Cytoscape^65^ and ClueGO^66^.

### Integrative scATAC-seq and scRNA-seq data analysis

The lists of more/less accessible regions in the scATAC-seq closest gene analysis determined by the *Tet2*^-/-^ HSC versus WT HSC pairwise testing were intersected with the lists of genes showing higher/lower expression in the *Tet2*^-/-^ HSC versus WT HSC scRNA-seq analysis. To find genes related to identified scATAC-seq/scRNA-seq targets, the Genemania database^24^ was used. Network of TFs (scATAC-seq analysis) and genes they regulate (scRNA-seq analysis) was built using information from the DoRothEA database^30^. Plots were made with with VennDiagram^72^, ggplot2^62^, Genemania^24^, DoRothEA^30^ and igraph^73^.

### Sample preparation for proteome analysis

Hoxb8-FL cells were cultured in RPMI 1640 media (Sigma), supplemented with 10% FBS (Gibco), 0.1% mercaptoethanol (Invitrogen), 1% penicillin-streptomycin (Sigma), 1% glutamine (Sigma), 1 μM estradiol and 5% FLT3L conditioned media from the B16-FL cell line. Cells were maintained in culture at concentrations of 10^5^-10^6^ cells/ml. Prior to proteomic analysis, Hoxb8-FL cells were resuspended in phosphate buffered saline (PBS). For experiments ‘Direct 10K’ and ‘Direct 15K’, cell lysis was performed using 2% sodium dodecyl sulfate (SDS) with subsequent boiling at 95 °C. Cell lysates were sonicated and dried using vacuum centrifugation. Samples were re-suspended in 100 mM TEAB. Reduction and alkylation of cysteine residues was performed by incubation with a final concentration of 5 mM tris-2-carboxyethyl phosphine (TCEP) at 60 °C for 30 min followed by final concentration 10 mM iodoacetamide (IAA) for 30 min at RT protected from light. Protein-level isobaric labeling was performed using TMT 10plex reagents (Thermo Scientific) in accordance with manufacturer’s protocol. 100% (w/v) trichloroacetic acid (TCA) was added to the sample mixture at a ratio of 1 to 4, followed by incubation for 10 min. The sample was centrifuged at 14,000 rpm and the resulting protein pellet was re-suspended in 100 mM TEAB buffer. Trypsin was added and proteins were digested overnight at 37°C. For experiments ‘10K Fract I’ and ‘10K Fract II’, cells in a volume of 20 µl of PBS were thawed on ice and 2 µl 1 M TEAB, 1 µl 2% sodium dodecyl sulfate (SDS), and 1 µl Halt Protease & Phosphatase inhibitor cocktail (pre-diluted 1:5 in water) was added. Cells were lysed by bath sonication for 5 min followed by 3 min incubation at 90 °C. Reduction and alkylation of cysteine residues was performed by incubation with 2 µl 50 mM TCEP at 40 °C for 30 min followed by 1 µl 200 mM IAA for 30 min at RT protected from light. 0.5 µg trypsin was added and proteins were digested overnight at RT. Peptide-level Isobaric labeling was performed using TMT 10plex reagents (Thermo Scientific) in accordance with manufacturer’s protocol. Following quenching of the reaction with 5% hydroxylamine, samples were combined and dried completely by vacuum centrifugation. High pH Reversed-Phase (RP) fractionation was performed with the Waters XBridge C18 column (2.1 x 150 mm, 3.5 μm, 120 Å) on a Dionex UltiMate 3000 HPLC system. Ammonium hydroxide at 0.1% v/v was used as mobile phase A and mobile phase B was set as 100% acetonitrile / 0.1% v/v ammonium hydroxide. The peptide mixture was reconstituted in 100 μl mobile phase A and subjected to gradient elution at 200 μl/min as follows: 5 minutes isocratic at 5% B, for 15 min gradient to 35% B, for 5 min gradient to 80% B, isocratic for 5 minutes and re-equilibration to 5% (B). The chromatogram was recorded at 215 and 280 nm and fractions were collected every minute. Fractions were dried completely by vacuum centrifugation and stored at -20 °C until further use.

For primary mouse samples, 10,000-30,000 cells from *Tet2*^-/-^ and WT mice were FACS-sorted into 0.1 mL PCR tubes containing 20 µl ice-cold PBS and processed as described above for the ‘10K Fract I’ and ‘10K Fract II’ experiments. 6, 5 and 8 fractions were finally subjected to LC-MS analysis for the ‘10K Fract I’, ‘10K Fract II’ and primary mouse samples, respectively.

### LC-MS/MS analysis

LC-MS/MS analysis was performed on a Dionex UltiMate 3000 UHPLC system coupled with an Orbitrap Lumos Mass Spectrometer (Thermo Scientific). Each peptide fraction was reconstituted in 10 μL 0.1% formic acid and 7 μl were loaded on the Acclaim PepMap 100, 100 μm × 2 cm C18, 5 μm, trapping column with the μlPickUp method at a flow rate of 10 μl/min. The samples were subjected to a multi-step gradient elution on an EASY-Spray (75 μm × 50 cm, 2 μm) C18 capillary column (Thermo Scientific) at 45 °C. Mobile phase A was 0.1% formic acid and mobile phase B was 80% acetonitrile / 0.1% formic acid. The gradient separation method at flow rate 300 nl/min was as follows: for 90 min gradient 5% to 38% B, for 10 min up to 95% B, for 5 min isocratic at 95% B, re-equilibration to 5% B in 5 min, for 10 min isocratic at 5% B. Precursor ions were selected with mass resolution of 120k, AGC 4×10^5^ and max IT 50 ms in the top speed mode within 3 sec. Peptides were isolated for HCD fragmentation with quadrupole isolation width 0.7 Th and 50k resolution. Collision energy was set at 38% with AGC 1×10^5^ and max IT 105 ms. Targeted precursors were dynamically excluded from further isolation and activation for 45 seconds with 7 ppm mass tolerance. For the ‘Direct 10K’ and ‘Direct 15K’ runs a 150 min 5% to 38% B gradient was used. For the ‘10K Fract II’ experiment, the 5 fractions were injected twice by setting a maximum intensity threshold at 5×10^6^ in the second run (from 5×10^20^).

### Protein identification and quantification

MS raw data was searched against the SwissProt mouse database (16,945 entries) using the SequestHT node in Proteome Discoverer 2.2. Precursor mass tolerance was 20 ppm and fragment ion mass tolerance was 0.02 Da. Spectra were searched for fully tryptic peptides with no more than 2 missed cleavages and a minimum length of 6 amino acids. TMT6plex at N-termini and lysine residues and carbamidomethyl at cysteine residues were set as fixed modifications. Methionine oxidation and glutamine and asparagine deamidation were set as dynamic modifications. Peptide FDR was set to 0.01 and validation was based on q-value and target-decoy database search using the Percolator node. The Reporter Ion Quantifier node included a custom TMT-10plex quantification method with an integration window tolerance of 15 ppm. At least one unique peptide was required for identification and only unique peptides were used for quantification.

### Bioinformatic analysis of proteomic and bulk RNA-seq data

Scaled quantitative values were obtained by dividing each TMT signal-to-noise (S/N) ratio by the mean TMT S/N across samples per protein. For bulk RNA-seq data, the gene list was filtered for genes with a minimum of 2 libraries with a minimum count per million (CPM) of 1. Remaining CPM values were normalized using the trimmed mean of M values (TMM) method in the edgeR R package (version 3.40.2). Principal component analysis (PCA) was performed using the R package PCAtools (version 2.10.0). The bottom 10% least variable genes/proteins were not included in PCA. Correlation between proteome and bulk transcriptome data was assessed using the Pearson correlation coefficient. For shortlisting targets for followup analysis, proteins with an absolute log2 FC of > 0.5 across all comparisons between the two *Tet2*^-/-^ and WT replicates were considered potential targets. For the bulk RNAseq dataset, a Students’ t test was performed and genes with an absolute log2 FC > 0.5 and adjusted p-value < 0.05 were considered potential targets. KEGG and Reactome pathway analysis were performed using the limma^74^ (version 3.54.2) and ReactomePA^75^ (version 1.42.0) package, respectively. Interaction network analysis was performed using the STRING database^76^ with a combined score cutoff of 0.4. Networks were visualized in Cytoscape^65^ (version 3.10.0).

### Statistical analysis

For all other experiments, differences between groups were assessed by one or two-tailed Students’ t test (two groups) or one-way ANOVA with Tukey’s post hoc test (three or more groups) using Prism software (GraphPad). Error bars represent SD. ∗∗∗∗p < 0.0001, ∗∗∗p < 0.001, ∗∗p < 0.01, and ∗p < 0.05 and ns= non-significant.

