## Supplementary figures and images for "Selection of pre-leukemic hematopoietic stem cells driven by distinct extracellular matrix molecules"

### Figure S1

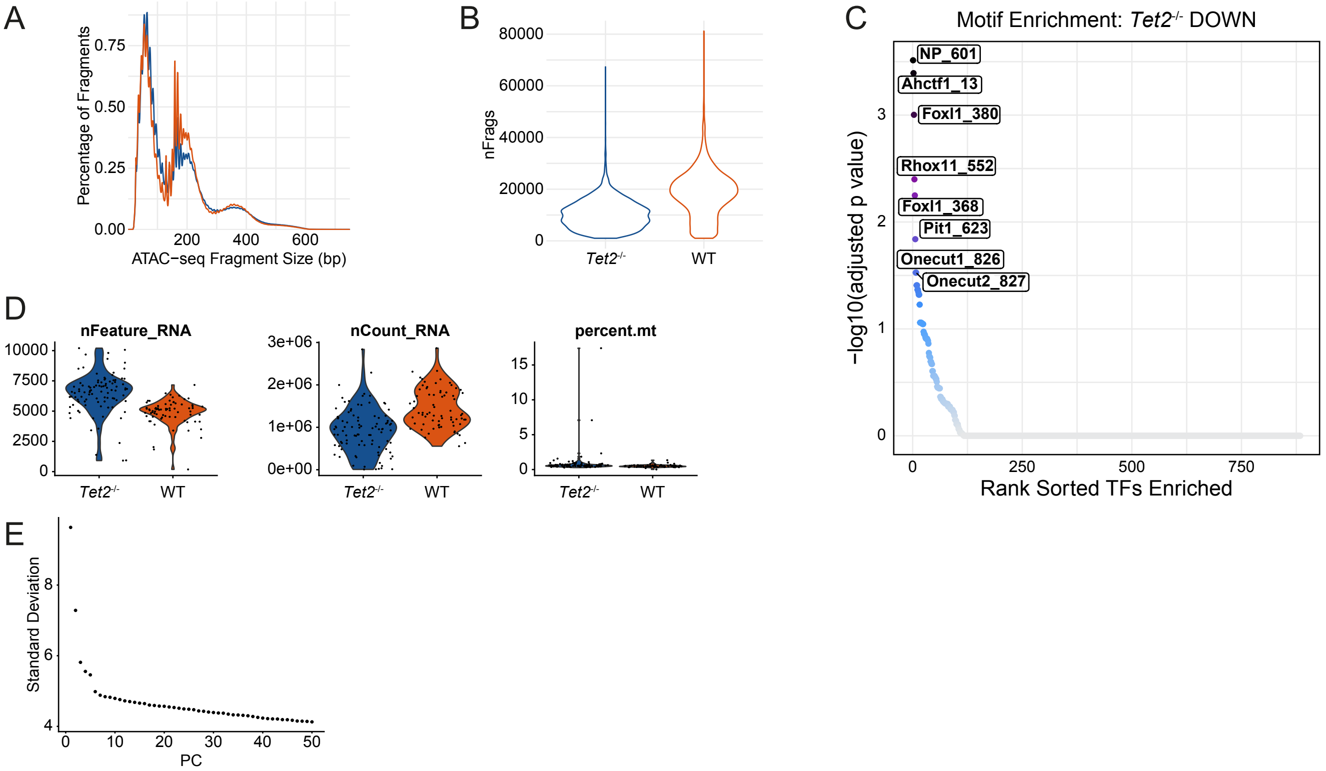

### Figure S2

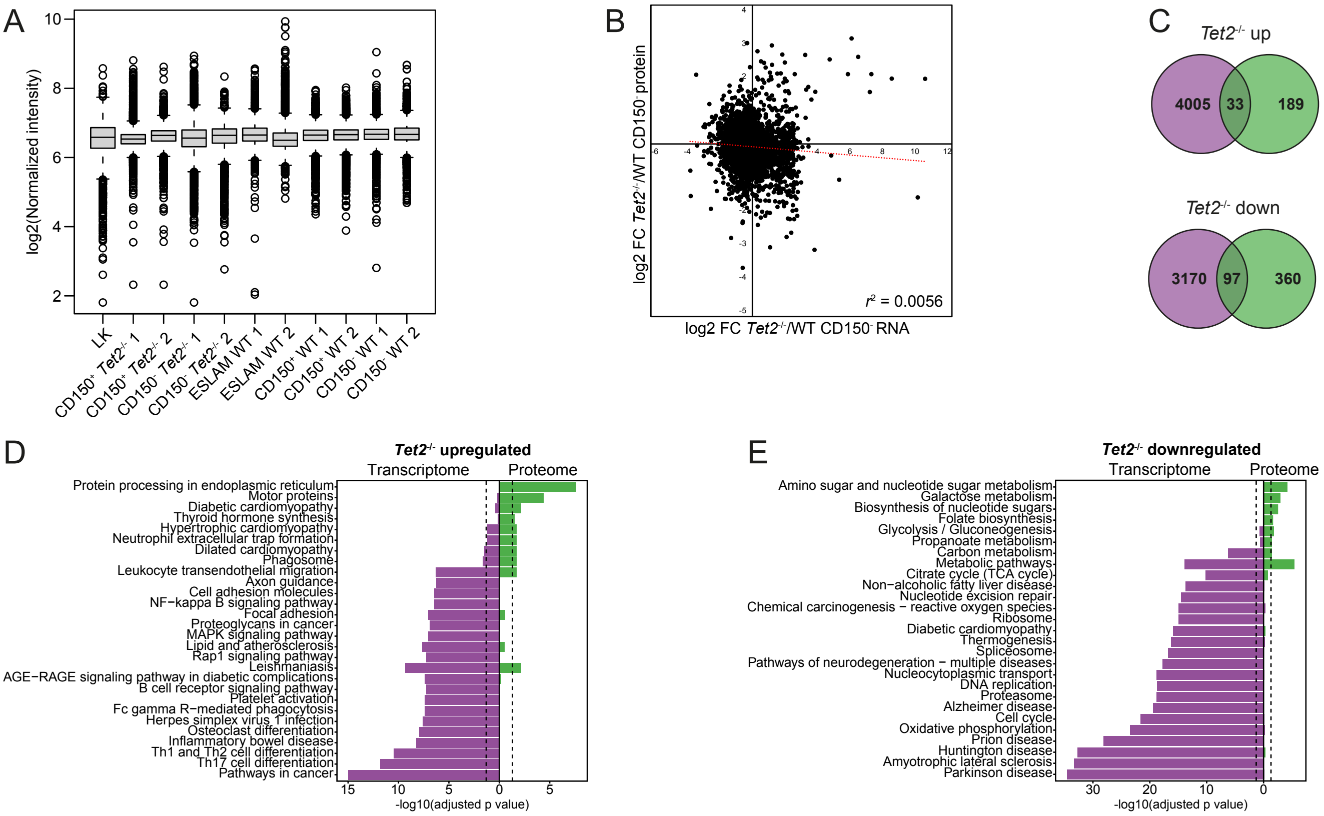

### Figure S3

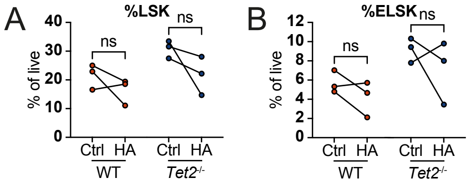
